# Human vascular organoids with a mosaic *AKT1* mutation recapitulate Proteus syndrome

**DOI:** 10.1101/2024.01.26.577324

**Authors:** Siyu He, Yuefei Zhu, Shradha Chauhan, Daniel Naveed Tavakol, Jong Ha Lee, Rayna Batya-Leia Berris, Cong Xu, Jounghyun H. Lee, Caleb Lee, Sarah Cai, Shannon McElroy, Gordana Vunjak-Novakovic, Raju Tomer, Elham Azizi, Bin Xu, Yeh-Hsing Lao, Kam W. Leong

**Affiliations:** Department of Biomedical Engineering, Columbia University, New York, NY 10027, USA; Irving Institute for Cancer Dynamics, Columbia University, New York, NY10027, USA; Department of Biological Sciences, Columbia University, New York, NY 10027, USA; Department of Computer Science, Columbia University, New York, NY 10027, USA; Data Science Institute, Columbia University, New York, NY 10027, USA; Herbert Irving Comprehensive Cancer Center, Columbia University, New York, NY 10032, USA; Department of Psychiatry, Columbia University Medical Center, New York, NY 10032, USA; Department of Pharmaceutical Sciences, University at Buffalo, The State University of New York, Buffalo NY 14214, USA; Department of Systems Biology, Columbia University Irving Medical Center, New York, NY 10032, USA; Center for Healthcare Innovation, Stevens Institute of Technology, Hoboken, NJ 07030, USA; Department of Medicine, Columbia University, New York, NY 10032, USA

## Abstract

Vascular malformation, a key clinical phenotype of Proteus syndrome, lacks effective models for pathophysiological study and drug development due to limited patient sample access. To bridge this gap, we built a human vascular organoid model replicating Proteus syndrome’s vasculature. Using CRISPR/Cas9 genome editing and gene overexpression, we created induced pluripotent stem cells (iPSCs) embodying the Proteus syndrome-specific AKT^E17K^ point mutation for organoid generation. Our findings revealed that AKT overactivation in these organoids resulted in smaller sizes yet increased vascular connectivity, although with less stable connections. This could be due to the significant vasculogenesis induced by AKT overactivation. This phenomenon likely stems from boosted vasculogenesis triggered by AKT overactivation, leading to increased vascular sprouting. Additionally, a notable increase in dysfunctional PDGFRβ + mural cells, impaired in matrix secretion, was observed in these AKT-overactivated organoids. The application of AKT inhibitors (ARQ092, AZD5363, or GDC0068) reversed the vascular malformations; the inhibitors’ effectiveness was directly linked to reduced connectivity in the organoids. In summary, our study introduces an innovative in vitro model combining organoid technology and gene editing to explore vascular pathophysiology in Proteus syndrome. This model not only simulates Proteus syndrome vasculature but also holds potential for mimicking vasculatures of other genetically driven diseases. It represents an advance in drug development for rare diseases, historically plagued by slow progress.

## Introduction

Proteus syndrome (PS), a rare genetic disorder, arises from a post-zygotic somatic mutation in the AKT gene (c.49G→A)^1^. This mutation, resulting in an amino acid change (AKT1^E17K^) within the AKT kinase’s Pleckstrin homology domain (PHD), leads to aberrant overactivation of the PI3K/AKT pathway. Characterized by tissue overgrowth, vascular malformations, cerebriform connective tissue nevi, subcutaneous tumors, and various skeletal abnormalities^2,3^, PS’s pathogenesis remains challenging to study due to its rarity (<1 case per million people) and diagnostic complexity^4,5,6^, often confused with similar syndromes like Cowden syndrome, which features different PTEN mutations^7,8^.

Limited by the availability of patient samples and disease models, PS drug development has been unsatisfactory. In vitro 2D culture systems with patient-derived cells can provide only limited pathophysiological information. Notably, Coriell Institute is the only depository that has the PS patient’s primary fibroblast and patient-derived lymphocyte line; yet, our sequencing results indicated that both did not carry AKT1^E17K^ (**Supplementary Fig. 1**). Chimera animal models^9^ with germline transmission of the conditional *AKT1* allele recapitulate the clinical manifestations of PS syndrome in the skin, bones, and vasculature, but the complexity and high variability of these models make them difficult to use for drug screening. Animal models are also unsuitable for identifying patient-specific treatment responses due to variations across species. These challenges have hindered the clinical management of PS patients^10^. Here, we propose a vascular organoid model to model the vascular manifestations of PS.

By introducing the AKT1^E17K^ mutation into wild-type induced pluripotent stem cells (iPSCs) via CRISPR gene editing, we built engineered vascular organoids with a mosaic mutation rate of 50% at the target locus, aligning with clinical observations^11^. In parallel, we also created a different set of PS-mimicking vascular organoids using iPSCs overexpressing AKT1^E17K^ or wild-type AKT1 (AKT1^WT^) for comparison. In both organoid models, we observed enhanced angiogenesis during the early organoid development. Decreased organoid size and an enhanced blood vessel network were found in the *AKT1*-mutated vascular organoids, consistent with the vascular malformation systematically seen in Proteus syndrome patients, who can have the single-channel form of the disease (capillary, lymphatic, or venous) or exhibit defects in multiple channel types^4,12^. We then generated a semi-3D vascular network to test three *AKT1* inhibitors. ARQ092 (miransertib) was the most effective in reducing the excessive complexity of the *AKT-activated* vasculature. These iPSC-derived organoid models recapitulated the clinical phenotypes of vascular malformation observed in PS patients and produced the expected drug response. They should facilitate drug discovery for PS treatment.

Traditional 2D in vitro models offer limited pathophysiological insights, while complex chimera animal models^9^ with germline transmission of the conditional *AKT1* allele though recapitulating PS clinical manifestations, prove unwieldy for drug screening and personalized response studies^10^. To overcome these limitations, we developed a vascular organoid model to emulate PS’s vascular features. Using CRISPR gene editing, we introduced the AKT1^E17K^ mutation into wild-type induced pluripotent stem cells (iPSCs), achieving a 50% mosaic mutation rate in the engineered vascular organoids, mirroring clinical observations^11^. A parallel organoid set with iPSCs overexpressing AKT1^E17K^ or wild-type AKT1 (AKT1^WT^) was also developed for comparative analysis.

In both organoid models, enhanced angiogenesis was observed during early development stages, along with reduced size and increased blood vessel networks, aligning with PS’s vascular abnormalities. These findings encompass both single-channel (capillary, lymphatic, or venous) and multi-channel vascular defects typical in PS. Subsequently, we assessed the efficacy of three AKT1 inhibitors on a semi-3D vascular network within these organoids. ARQ092 (miransertib) demonstrated notable success in moderating the heightened vascular complexity induced by AKT activation. This iPSC-derived organoid approach not only replicates PS’s clinical vascular malformations but also aligns with expected drug responses, offering a promising avenue for PS treatment advancements. These models provide valuable tools for furthering drug discovery in treating this rare and complex syndrome.

## Results

### Generation of iPSC-derived vascular organoids with the *AKT1*^E17K^ mutation

#### Obtaining iPSCs carrying mosaic AKT1^E17K^

To generate iPSCs carrying the *AKT1*^E17K^ mutation, we applied CRISPR/Cas9 gene editing followed by pool selection (**Fig. 1a, Methods**). We first nucleofected wild-type iPSCs with the high-fidelity *Streptococcus pyogenes* Cas9 endonuclease, a guide targeting the mutation site and a single-stranded homology-directed repair (HDR) template. Initial editing efficiency reached 17.6% with a non-PAM HDR template, compared to less than 1% for the PAM-containing template (**Fig. 1b**), likely due to Cas9-mediated cleavage. The nucleofected iPSCs were then sorted into 96-well plates at a density of 10 cells per well. Notably, significant variability in HDR efficiency was observed across the 44 sorted wells (**Supplementary Fig. 2a**). To optimize the G→A substitution rate at the target locus, we used limiting dilution cloning (LDC) on the selected pool. After four rounds of selection, the G→A ratio increased to 38.9% (**Supplementary Fig. 2b**). Sanger sequencing after the fifth round revealed that 50% of sequences from the target site exhibited the guanine to adenine change (**Fig. 1c**). Due to reports of the homozygous c.49G→A mutation being lethal^13^, we concluded selection, hypothesizing that the final iPSC pool might consist of clones heterozygous for the c.49G→A mutation. Whole-genome sequencing of the fifth-round cells confirmed our hypothesis, showing a G→A substitution rate of 28% (**Supplementary Fig. 2**), aligning with the mosaic mutation rate of 1-50% observed in Proteus syndrome patients^1^. Immunocytochemistry results further validated the predominant expression of AKT1^E17K^ in the cells from our final pool (**Supplementary Fig. 2c**). These findings demonstrate the successful creation of iPSCs with the AKT1^E17K^ mutation desired for subsequent organoid studies.

**Figure 1.**
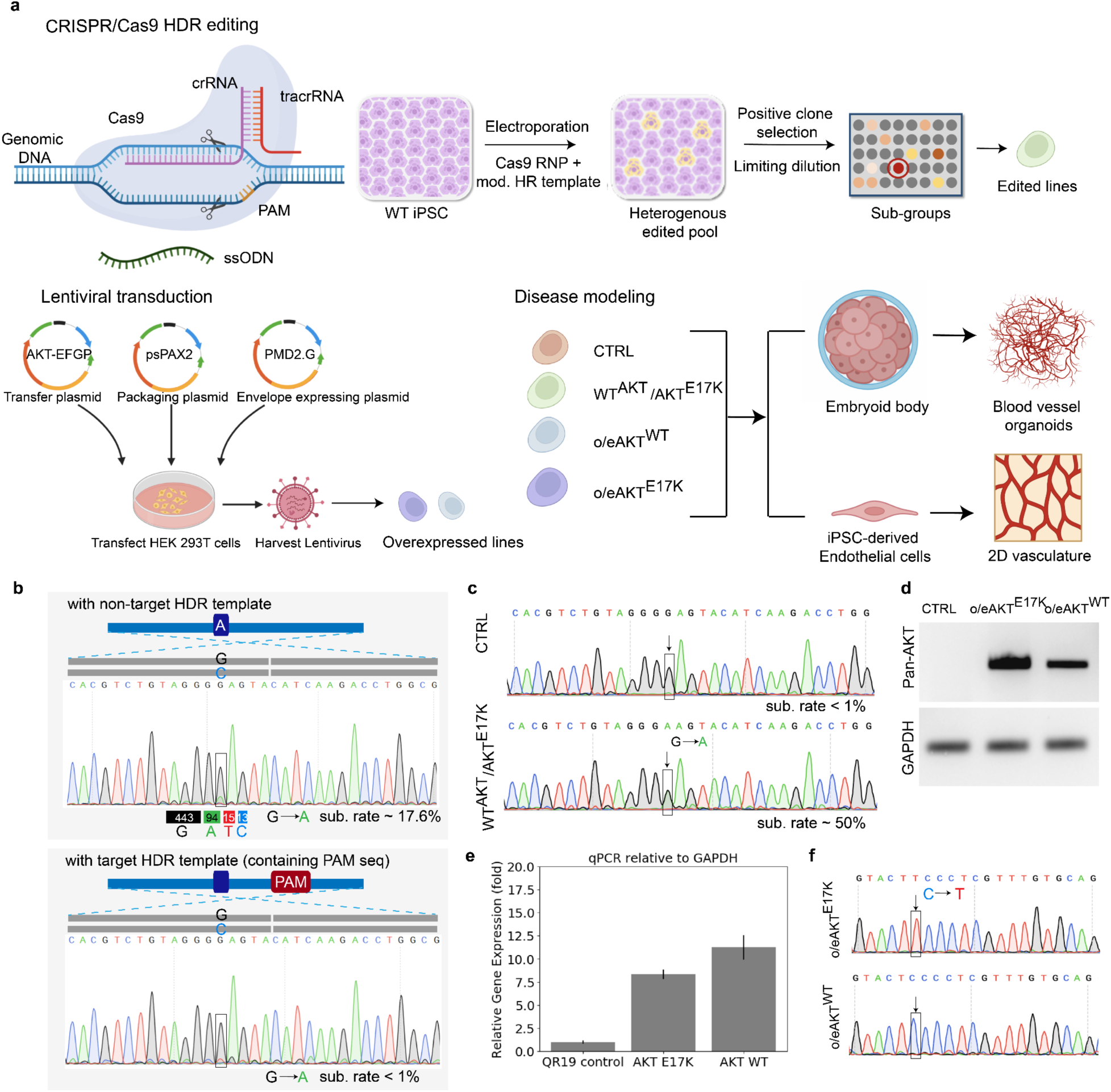
CRISPR editing and lentiviral transduction to generate Proteus syndrome iPSCs. (a) Processes of CRISPR/Cas9 editing and lentiviral transduction used to generate AKT^E17K^-overactivated iPSCs. (b) Editing efficiencies comparing ‘nontarget’ and ‘target’ HDR templates in iPSCs. (c) Sanger sequencing of final iPSC pool after selection. (d) Gel electrophoresis validating AKT overexpression in transduced iPSCs. (e) Quantification of the AKT1 level in AKT-overexpressing iPSCs. (f) Sanger sequencing confirming AKT1 mutation in overexpressed iPSCs.

#### Obtaining iPSCs overexpressing AKT1

We introduced EGFP-tagged wild-type AKT1 or AKT1^E17K^ into iPSCs (**Supplementary Fig. 2d, Methods**). RT‒PCR analysis confirmed the overexpression of AKT1 in AKT^E17K^ (o/eAKT^E17K^) and AKT^WT^ (o/eAKT^WT^) cell lines compared to nontransduced controls (**Fig. 1d**). RT‒ qPCR results were consistent, showing *AKT1* overexpression in both cell lines: an 8.3-fold increase in o/eAKT^E17K^ and an 11.2-fold increase in o/eAKT^WT^ compared to parental controls (**Fig. 1e**). Sanger sequencing further verified the *AKT1* overexpression in o/eAKTE17K cells (**Fig. 1f**).

#### Characterization of Cas9-edited and AKT1-overexpressing iPSCs

AKT activation, achieved in Cas9-edited (AKT^E17K^) and AKT1-overexpressing (o/eAKT^E17K^ and o/eAKT^WT^) iPSCs, downregulated key AKT pathway genes, including *VEGFA*^14^, *ZDHHC8*^15^, and *PIK3CA*^16^, indicating successful modulation of *AKT1* activity through either gene editing or overexpression approaches. In contrast, VEGFR expression increased in overactivated AKT^E17K^ and o/eAKT^E17K^ cells (1.57-fold higher than control) but decreased in o/eAKT^WT^ cells (65% lower than control), indicating divergent effects of AKT1 activation and wild-type overexpression on VEGFR expression (**Supplementary Fig. 4**).

#### Generation of blood vessel organoids

To model the vascular malformations seen in PS, we generated blood vessel organoids using previously established protocols^17,18^ (**Fig. 2a, Methods)**. The organoids were fixed for light sheet microscopic imaging on day 11^19^. Staining for endothelial (CD31) and mural (PDGFRβ) cell markers confirmed the formation of a vascular network composed of endothelial tubes and mural cells, allowing for the spatial visualization of these cells within the organoid vasculature (**Fig. 2b**).

**Figure 2.**
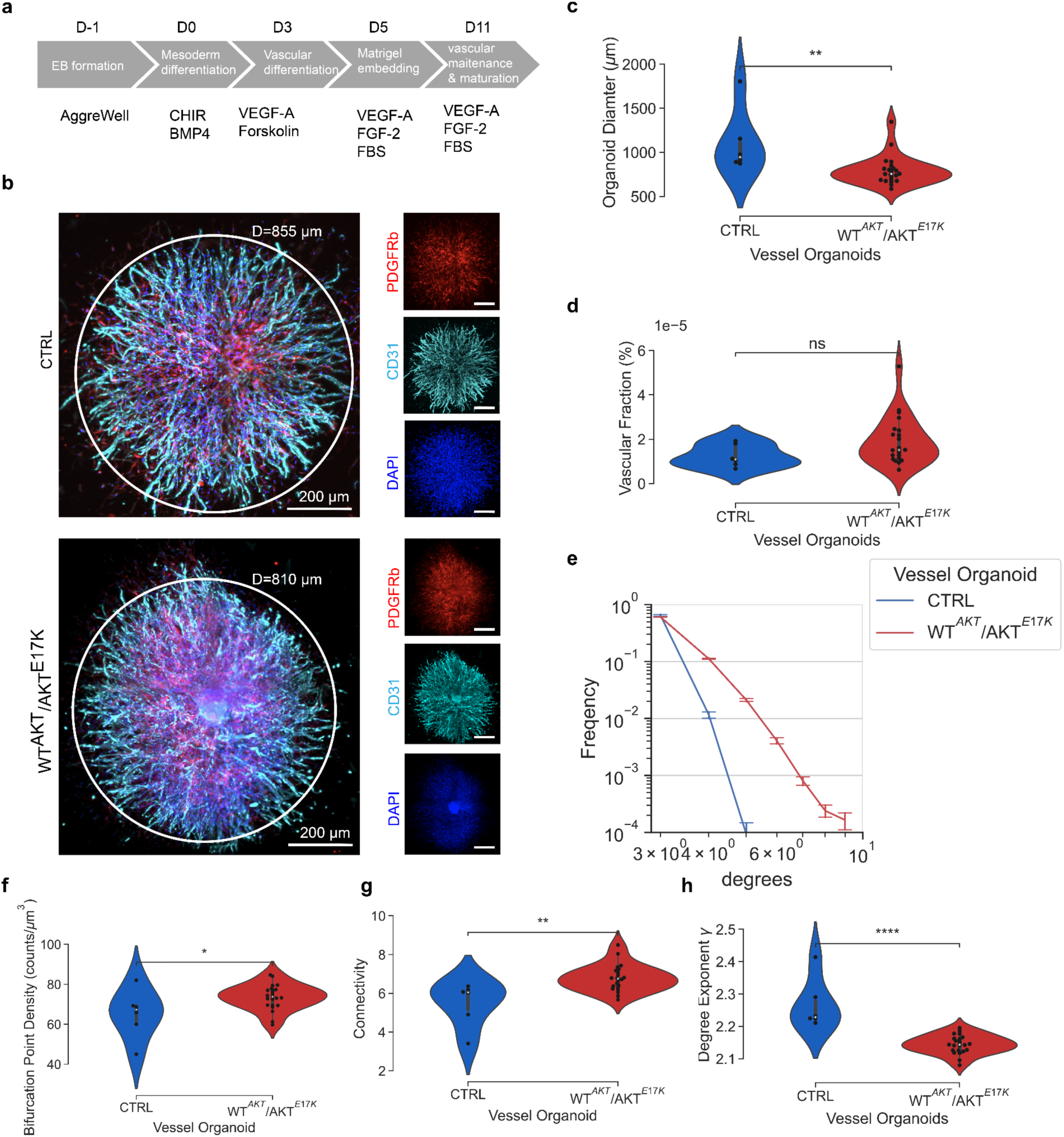
Generation and characterization of AKT-overactivated vascular organoids. (a) Step-by-step protocol for blood vessel organoid formation from iPSCs. (b) Light sheet microscopy images of vascular organoids from both wild-type and AKTE17K gene-edited iPSCs, stained with CD31, PDGFRβ, and DAPI. Insets highlight individual channel fluorescence. (c-h) Comparative analysis of AKT^E17K^ and control organoids, examining organoid size, vascular fraction, network degree distribution, bifurcation density, connectivity, and degree exponents. Statistical analysis: one-tailed Student t-test; significance denoted as n.s. (no significance), * (p≤0.05), ** (p≤0.01), *** (p≤0.001), **** (p≤0.0001); n=6 for control organoids, n=22 for AKT^E17K^ organoids.

### AKT overactivation induced the overgrowth of vascular phenotypes

Investigating the defects in 3D vasculature induced by AKTE17K mutation presented a challenge due to the complexity of the vessel structures (**Fig. 2b**). To overcome this, we applied our previously developed machine learning-based computational methods^20^. This involved segmenting vessel tubes and reconstructing network structures and graphs for each AKT^E17K^ iPSC-derived blood vessel organoid, summarizing their phenotypical and topological features (see **Methods**). Notably, despite initial size standardization at the embryoid body (EB) stage (**Fig. 3a, Supplementary Fig. 5a**), AKT^E17K^ organoids were significantly smaller in size (**Fig. 2c**). The vascular fraction, based on density estimation of segmented vessel organoids (**Fig. 2d**), showed no significant difference, indicating comparable vascular density between mutated and control groups.

**Figure 3.**
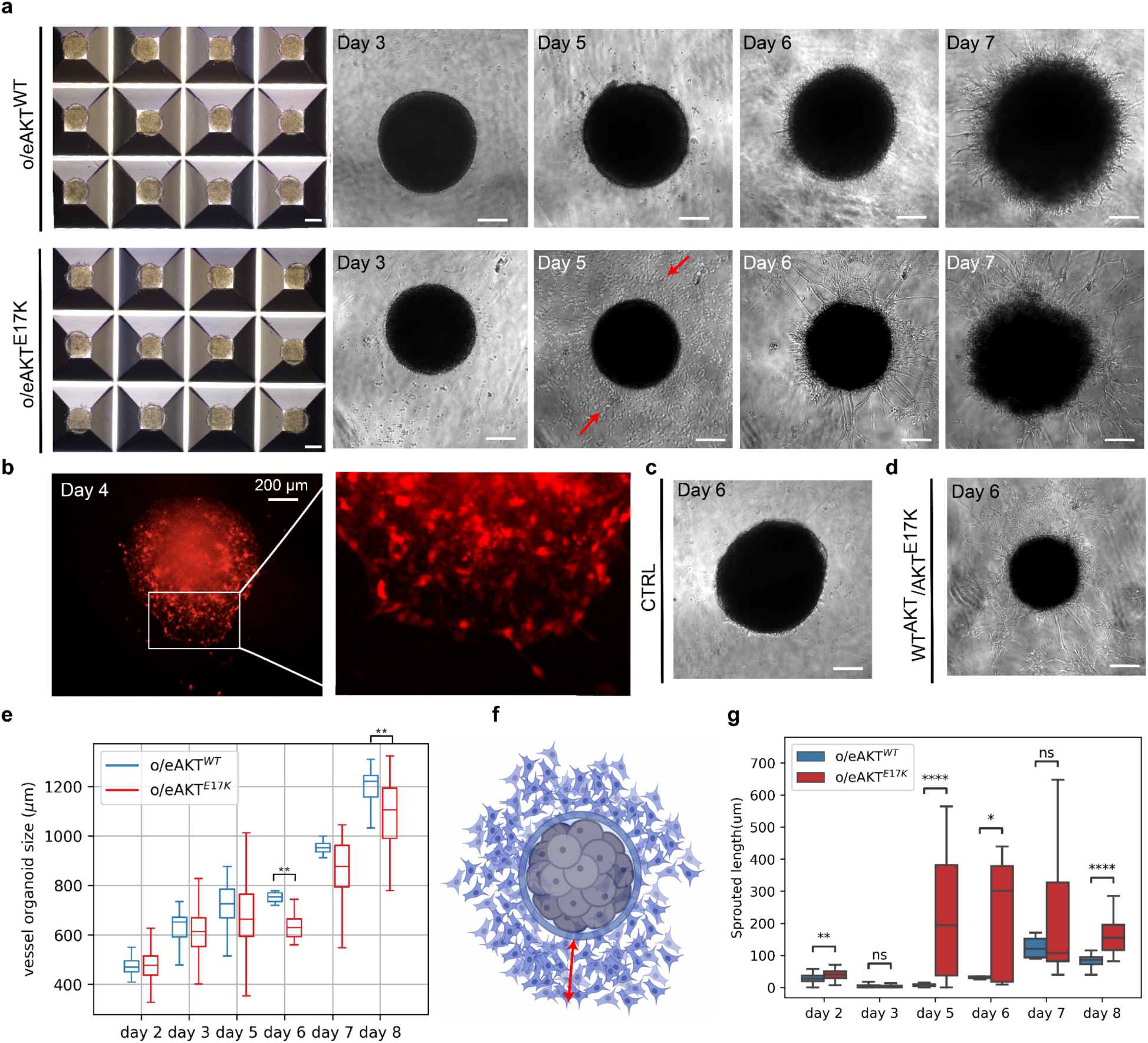
Enhanced Vasculogenesis in *AKT*-Overactivated Organoids. (a) Representative bright field images of vascular organoids derived from o/eAKT^WT^ and o/eAKT^E17K^ iPSC lines on days 1, 3, 5, 6, and 7 (scale bar: 200 μm). (b) Representative fluorescence images of mCherry-labeled AKT^E17K^ organoids on day 4. Representative bright-field images of (c) the control and (d) AKT^E17K^ organoids on day 6. (e) Comparison of size between o/eAKT^WT^ and o/eAKT^E17K^ iPSC-derived organoids, analyzed using one-tailed, unpaired Student’s t-test. (f) Scheme of cell migration on the vascular organoids on day 5. The red arrow denotes the migration distance between the sprouting cells and the organoid core. (g) Quantification of the distance of cell migration for o/eAKT^WT^ and o/eAKT^E17K^ iPSC-derived organoids, compared using one-tailed Student’s t-test. Significance is presented as follows: n.s., no significance; *, p<=0.05; **, p<=0.01; ***, p<=0.001; ****, p<=0.0001.

For a deeper understanding of vascular network connectivity, we analyzed the organoids using network graphs, treating bifurcation points and end points of vascular branching as nodes, and branches as edges. The degree of each bifurcation point was defined by the number of connections to other nodes. Compared with that of the wild-type organoids, the degree of the vascular network in the AKT^E17K^ organoids always had a higher frequency (**Fig. 2e**) and a higher bifurcation point density (**Fig. 2f**). The features of these two variables resulted in higher connectivity (**Fig. 2g**), defined as the endpoint density divided by bifurcation points. Moreover, the degree exponent of AKT^E17K^ organoids had a lower value than that of the wild-type control, indicating less stable connections within the vascular network of the diseased organoids (**Methods**).

### Significant vasculogenesis occurred in *AKT*^E17K^ vascular organoids at early time points

To investigate whether the *AKT*^E17K^ mutation affected early vascular development, we traced the morphological development of the organoids with phase contrast images. Instead of embedding the organoids in Matrigel®, we placed the organoids on top of the gel for imaging. By day 5 (two days after vascular induction), we noticed that the cells of the *AKT*^E17K^ organoids migrated to the surrounding periphery (shown with red arrows). By day 6 (one day after Matrigel embedding), rapid formation and expansion of vascular tubes were observed in these organoids (**Fig. 3a**). In contrast, the control and wild-type AKT1-overexpressing (o/eAKT^WT^) groups exhibited no cell sprouting on day 5 and minimal vascular sprouting on day 6, although slightly more developed sprouting appeared by day 7. mCherry-labeled cells revealed budding at the boundaries of AKTWT organoids by day 4 (**Fig. 3b-d, Supplementary Fig. 5b**), but absent in control organoids (**Supplementary Fig. 5b**). In bright field images, the AKT^E17K^ organoids were visibly smaller than their control counterparts (**Fig. 3c, 3d)**.

We measured the organoid size (excluding sprouting areas) in both o/eAKT^WT^ and o/eAKT^E17K^ groups throughout their development (**Fig. 3e**). From day 3 to day 6, a slowed growth rate was observed, likely due to mesodermal progenitors differentiating into vascular lineage. Post day 6, the growth rate increased with vascular structure formation. The size difference, particularly on days 6 and 8, indicated that o/eAKT^E17K^ organoids were consistently smaller, as also seen in light sheet microscopic imaging (**Fig. 3e**).

Further assessment of sprouting cells and vessel tubes revealed significantly longer vessel lengths in o/eAKT^E17K^ organoids compared to o/eAKT^WT^, ranging from approximately 100 to 500 µm (**Fig. 3f, g**). By day 11, well-formed vascular networks were evident, as shown by CD31 staining (**Supplementary Fig. 5c**). These results suggest that the AKT^E17K^ mutation not only enhances vascular sprouting but also promotes vascular structure formation, a phenotype corroborated by other in vivo studies^21^.

### Perivascular cell dysfunction in AKT-overactivated organoids

To investigate the dynamics of cell composition in our vascular organoids, we quantified the signal areas of PDGFRβ and CD31 from the confocal images at the pixel level as the representative relative cell composition (**Fig. 4a, Methods**). As Matrigel itself might interfere with the imaging of organoids, we removed the gel prior to confocal imaging; however, this might disrupt the organoid architecture and shrink the vascular tubes. Thus, the confocal images of our blood vessel organoids might look different than those obtained via light sheet microscopic imaging. In the AKT-overactivated organoids (o/eAKT^E17K^ and AKT^E17K^), we found significantly more PDGFRβ+ mural cells (**Fig. 4b, 4c**). To further determine whether these mural cells were dysfunctional in the AKT-overactivated samples, we generated vascular organoids without gel embedding. This approach excluded the exogenous extracellular matrix, focusing instead on the matrix produced by the mural cells (including smooth muscle cells and pericytes) to support vascular tube formation. Surprisingly, control iPSC-derived organoids formed vascular tubes even without Matrigel, although with a sparser network (**Fig. 4d**). In contrast, AKT^E17K^ organoids lacked CD31 signals, displaying only PDGFRβ. We inferred that mural cells in AKT-overactivated organoids were dysfunctional in matrix secretion.

**Figure 4.**
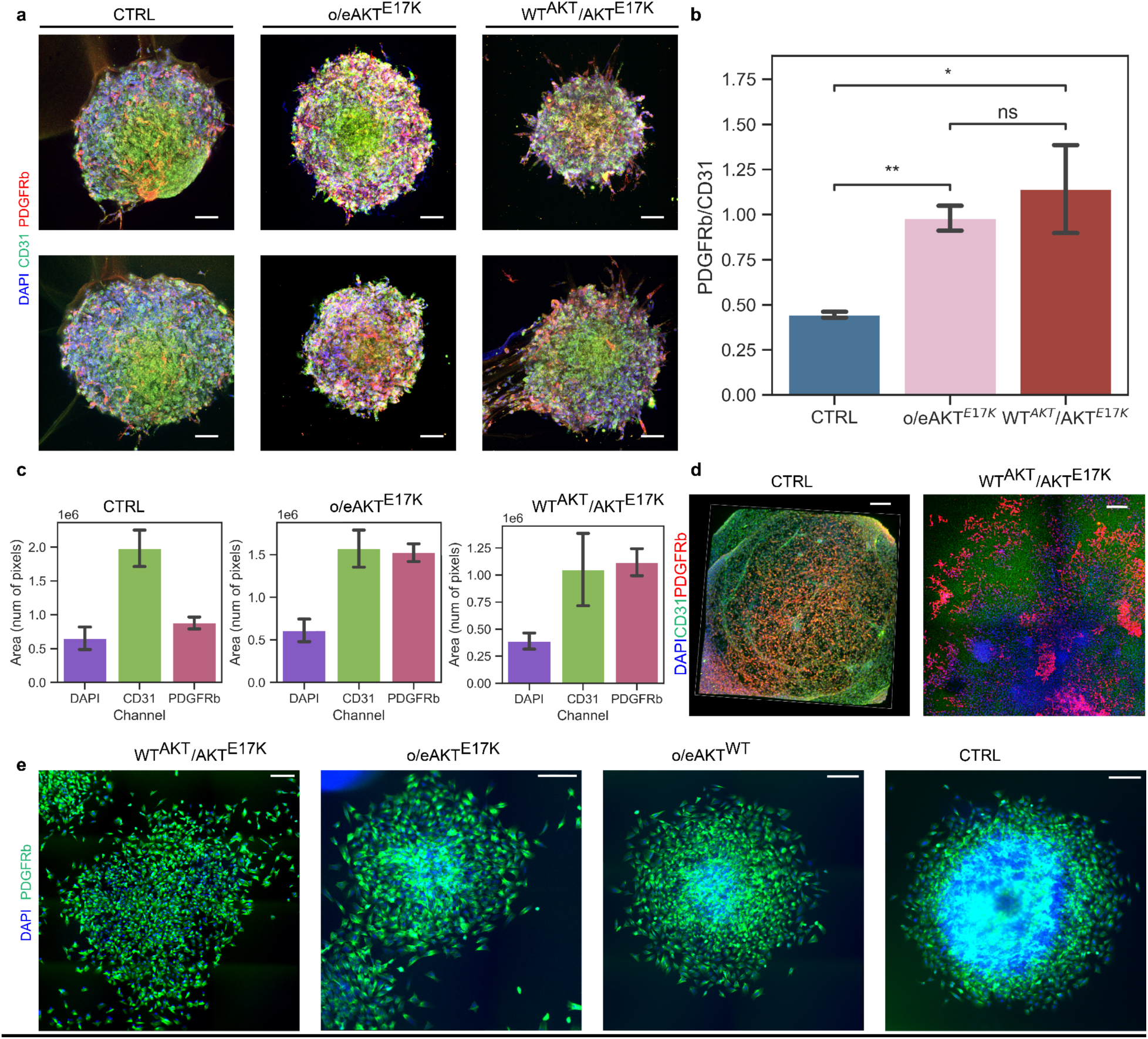
Perivascular cell dysfunction in AKT-overactivated organoids. (a) Representative confocal images of organoids stained with a CD31 antibody, a PDGFRβ antibody, and DAPI (scale bar: 200 μm). (b) Signal ratio of PDGFRβ to CD31 in different organoids. A one-tailed t test was performed to compare the differences between groups. Significance is presented as follows: n.s., no significance; *, p<=0.05; **, p<=0.01. (c) Calculated signal areas in confocal images (unit: pixel). (d) Representative confocal images of vascular organoids cultured without Matrigel (scale bar: 200 μ m). (e) Representative fluorescence images of SMC spheroids generated with o/eAKT^E17K^, o/eAKT^WT^, AKT^E17K^, and control iPSCs (stained with a PDGFRβ antibody and DAPI).

Confirming this dysfunction, we differentiated iPSCs into smooth muscle cells (SMCs) and generated SMC spheroids marked by PDGFRβ expression (**Fig. 4e**). However, the AKT^E17K^ SMC spheroids failed to maintain their shape, with cells dispersing outward, unlike the control groups that retained spheroidal form. This finding suggests that the mural cells in AKT-overactivated conditions exhibit matrix secretion dysfunction, evident from their inability to form stable spheroids.

### AKT inhibitors rescued vascular malformation driven by AKT overactivation

To assess the potential of vascular organoids as a drug testing platform for Proteus syndrome, we treated them with three AKT inhibitors: Miransertib (ARQ092), Capivasertib (AZD5363), and Ipatasertib (GDC0068). ARQ092, an FDA-approved AKT inhibitor for PS^23, 24^, has shown efficacy in reducing facial bone overgrowth and the size of cerebriform connective tissue nevi in PS patients after one year of treatment, although a survey of 41 patients indicated no significant improvement in overall clinical outcomes as per the Clinical Gestalt Assessment (CGA) for PS^10^. Both AZD5363 and GDC0068, still under preclinical evaluation for tumor treatment^27, 28^, are known for their anti-metastasis effects^29^. Our organoid approach offers a potentially more robust and clinically relevant in vitro model for drug discovery, given its high throughput and ability to mimic human-specific phenotypes.

To simplify the comparison of drug responses, we bypassed the effect of the extracellular matrix and dysfunctional mural cells on organoids by dissociating the 3D vascular organoids into single cells and placing them on Matrigel to form 2D vascular networks^30^ (**Fig. 5a, Methods**). We then treated these networks with 1 μM of each AKT inhibitor, a concentration previously reported as effective^24^ (**Fig. 5b**). To quantify the drug effect on network morphology, we applied a deep learning method^31^ to segment the vessel network and extracted the skeleton from the segmentation to generate the 2D network graph (**Fig. 5c**). When compared with that of the parental control, the network connectivity was significantly increased in the AKT^E17K^ organoids (**Fig. 5d**), which again confirmed the effect of AKT overactivation. Treatment with AKT inhibitors significantly reduced this increased connectivity, with the extent of reduction (ARQ092>AZD5363>GDC0068) correlating with their known AKT inhibitory activities (IC50 values for AKT1: 2.7 nM, 3 nM, and 5 nM for ARQ092, AZD5363, and GDC0068, respectively^27,28,32,33^). Interestingly, o/eAKT^E17K^ organoids did not respond to the drug treatment. This could be due to unknown effects caused by lentivirus-induced random insertion of the AKT^E17K^ expression cassette into the genome. This result highlights the benefit of using the CRISPR/Cas9 technique for precise genome editing. Taken together, these results demonstrate the advantages of using engineered vascular organoids for modeling genetic vascular diseases and screening potential therapeutics.

**Figure 5.**
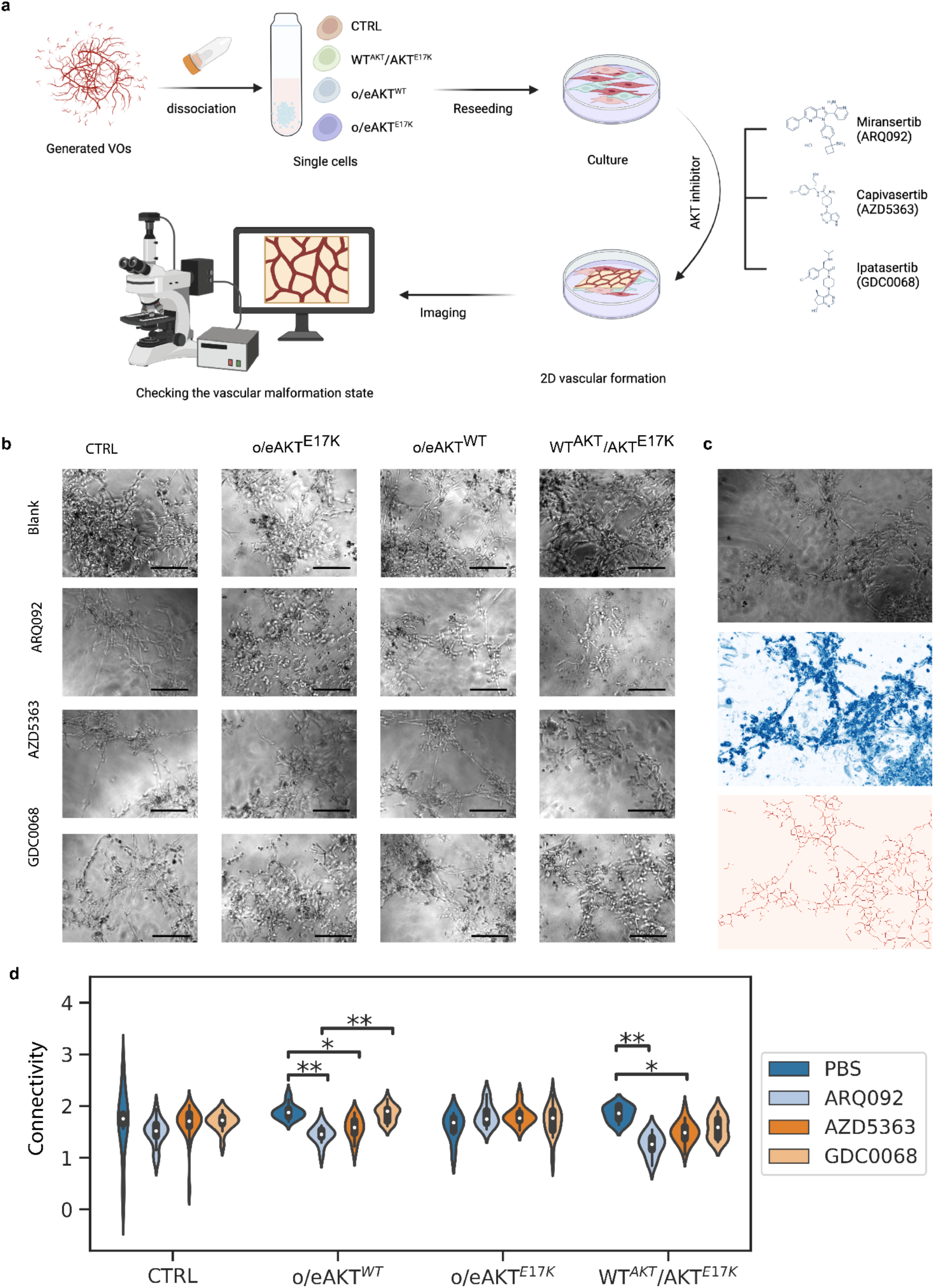
Effects of AKT inhibitors on vascular networks in AKT-overactivated samples. (a) Scheme of drug treatment in this study. Vascular organoids were dissociated into single cells and placed onto Matrigel-coated 96-well plates, followed by drug treatment. (b) Representative bright field images of blood vessel networks (scale bar: 200 μm). (c) Representative images showing segmentation and skeletonization of the network graph. (d) Graph showing the connectivity of vascular networks formed by control, o/eAKT^E17K^, o/eAKT^E17K^, and WT^AKT^/AKT^E17K^ ECs treated with ARQ092, AZD5363, and GDC0068. Two-way ANOVA was performed with the post hoc test. The p values obtained via ANOVA for iPSC type, treatment, and interaction were statistically significant (p=1.06e-05, 4.15e-09, 1.26e-06, respectively). Significance is presented as follows: n.s., no significance (not shown in the figure); *, p<=0.05; **, p<=0.01.

## Discussion

This study highlights the effectiveness of iPSC-derived organoids in disease modeling and drug screening, particularly for rare genetic disorders like Proteus Syndrome (PS), where patient sample scarcity and the limitations of animal and 2D cellular models pose significant challenges. Our use of organoids not only bypasses these obstacles but also facilitates the replication of patient-specific phenotypes and offers predictive insights into individual drug responses, thereby potentially reducing clinical trial risks.

Using CRISPR/Cas9 genome editing, we developed vascular organoids with the AKT^E17K^ mutation, mirroring the vascular malformation phenotype commonly observed in PS patients. These malformations, which include vascular hamartoma and port wine stains, occur in over 70% of PS patients^34^. Our model revealed that AKT overactivation leads to smaller but more interconnected vascular organoids with less stability (**Fig. 2c-h**), effectively simulating venous anomalies found in about 20-30% of PS patients. This is likely due to enhanced vasculogenesis triggered by AKT overactivation. Importantly, the homozygous AKT^E17K^ mutation is lethal, making the overexpression models (o/eAKT^WT^ and o/eAKT^E17K^) crucial for studying its impact on vascular formation. Consistent with in vivo studies, the AKT^E17K^ mutation in our model promoted vascular sprouting and vasculogenesis, especially in the o/eAKT^E17K^ organoids (**Fig. 3**).

We also observed a significant difference in cell composition between wild-type and diseased organoids. AKT overactivation led to an increased number of dysfunctional PDGFRβ+ mural cells in terms of matrix secretion (**Fig. 4a-c**), a finding further validated using iPSC-differentiated smooth muscle cells (SMCs) (**Fig. 4d**). This underlines the capability of our organoids to replicate both the disease and its vascular pathophysiology.

Treatment with AKT inhibitors, particularly ARQ092, demonstrated the potential to reverse vascular malformations in these organoids, highlighting their utility in drug screening. ARQ092’s efficacy in reducing vascular connectivity confirms the link between vascular dysfunction in PS and AKT overactivation. Future studies could expand on these findings by examining phenotypic variations based on allele frequency in organoids and comparing the effects of different inhibitors, such as mTOR and PIK3CA, with those of AKT inhibitors.

## Methods

### Enhanced CRISPR/Cas9 system for AKT1 E17K knock-in editing

A 20-nt guide sequence in crRNA was designed to target a specific region of the AKT gene. Guide RNAs and trans-activating crRNA were annealed and fused with the Cas9 protein to form ribonucleoprotein complexes (RNPs). Upon intracellular delivery by electroporation, the Cas9 endonuclease produced a double-strand break (DSB) at the site 3 nt downstream of the PAM sequence. The DSB caused nonhomology end joining (NHEJ) and homology-directed repair (HDR). HDR was enhanced by a PAM nontargeted single-stranded donor oligonucleotide (ssODN). To confirm the gene mutation, DNA was extracted, and the mutation rate was calculated by Sanger sequencing with peak height quantification.

### Lentiviral transduction for AKT1 overexpression

We followed our established protocols to generate AKT^WT^ and AKT^E17K^ overexpression lentiviral vectors^35,36^. Briefly, the AKT^WT^ or AKT^E17K^ lentiviral transfer plasmid (Addgene) was cotransfected into HEK293T cells (Takara) with the packaging plasmid psPAX2 (Addgene) and the envelope-expressing plasmid PMD2.G (Addgene) using the transfection reagent CalFectin (SignaGen) following the manufacturer’s recommendations. At 24 h posttransfection, the CalFectin-containing media were replaced with fresh complete media. The viral particle-containing media were collected at 48 and 72 h posttransfection. The collected media were then filtered with a 0.45 mm filter (VWR) to remove the cell debris, followed by concentration using an AMICON 100K MWCO filter. The viral titer was determined by RT‒qPCR using the titer kit from Applied Biological Materials prior to use.

### Generation of blood vessel organoids

To model the vascular malformations seen in PS, we generated blood vessel organoids using previously established protocols^17,18^ (**Fig. 2a, Methods)**. The embryoid body (EB) was first formed in Aggrewell® from iPSCs obtained one day prior to the induction of differentiation (Day −1). On day 0, CHIR and BMP4 were added to induce the differentiation of embryoids into the mesoderm germ layer. On day 3, vascular differentiation was induced in the organoids by applying VEGFA and forskolin. The differentiated organoids were then embedded in Matrigel® on day 5 and cultured in medium containing VEGFA, FGF2, and FBS. The organoids were fixed on day 11 and stained for light sheet microscopic imaging^19^. We stained for endothelial and mural cell markers (CD31^+^, PDGFRβ+, α SMA+, respectively) to confirm the generation of a vascular network constructed by endothelial tubes and mural cells, with visualization of the spatial arrangement of these cells in the vasculature (**Fig. 2b**).

### Light sheet microscopy images and deep learning analysis

Vascular organoids were processed with RapiClear (SunJin Lab, RC149001) to clear the tissue, mounted into quartz cuvettes and imaged by CLARITY-optimized light sheet microscopy^37^. Preprocessing of raw images included homomorphic filtering and contrast-limited adaptive histogram equalization before segmentation. Following the analysis in our previous study^20^, we applied VesSAP^38^ to CD31^+^ images. Skeletonization with the Python library “skimage” was performed on the segmented images. Skan and Networkx, two Python libraries, were then used to perform the network analysis of the vascular organoids. The vascular network graphs comprised the bifurcation points, which were defined as degrees, of the networks, *k*. The of the degree exponent was represented as parameters (*γ*) in the degree distribution *P*(*k*)∼*k*^-*γ*^.

### Confocal imaging

The blood vessel organoids were permeabilized with blocking buffer (1.5 ml FBS, 0.5 g of BSA, 250 μL of Triton X-100, 250 μL Tween 20 and 47.5 mL of PBS in 50 mL buffer) and incubated with primary antibody overnight at 4℃. The samples were then washed with PBS-T solution and embedded in mounting medium. A confocal microscope with a 10X objective was used to image the vascular organoids.

### Generation of SMC spheroids

SMC spheroids were generated via EBs using an established method to derive SMCs from iPSCs^22^. Briefly, one day after EB formation, the medium was changed to DMEM/F-12 containing N2, B27, CHIR-99021, and hBMP4. After 3 days of incubation, the medium was replaced with medium containing the growth factors Activin A and PDGF-ββ. The same medium was freshly made and added the next day. The SMC spheroids were then maintained in DMEM/F-12 containing Activin A and heparin.

### Drug treatment

Blood vessel organoids were dissociated into single cells with Accutase and collagenase on day 11. The cells were then seeded on 96 well Matrigel-coated plates by adding 100 µl (1.5 x 10^4^ cells) of the single-cell suspension. Vascular networks formed within hours. Then, 1 μM of Miransertib, Capivasertib, or Ipatasertib was added to the medium to treat the cells, and the vascular network was fixed for image analysis after two days of incubation.

## Author contributions

K. W. L. and Y. L. provided overall supervision of the study. S. H. and Y. L. performed CRISPR/Cas9 editing, lentiviral transfection, and whole-genome sequencing. S. H., Y. Z., D. N. T., J. H. L., R. B. B., and C. X. performed blood vessel organoid generation. S. H. performed data analysis and drug testing. S. H. and S. M. performed qPCR. S.C. and R.T. performed the light sheet microscopic imaging. S. H., Y. Z, D. N. T., J. H. L., R. B. B., C. X., J. H. L., C. L., S. C., S M, G. V. N., E. A., B. X., Y. L., and K. W. L. interpreted the data and wrote the manuscript. All authors reviewed, contributed to, and approved the manuscript.

## Notes

The authors declare no financial competing interests.

## Acknowledgments

This work was supported by NIH (UH3TR002151 to K.W.L.; R21NS133635 to Y.-H.L.), and Sigma Xi Society GIAR Grat (G03152021116627040 to S.H.). These studies used the resources of the Herbert Irving Comprehensive Cancer Center Confocal and Specialized Microscopy Shared Resource, funded in part through NIH/NCI Cancer Center Support Grant P30CA013696.

## Data and code availability

The data and code supporting the findings of this study will be released via the GitHub repository.

## Supplementary figures

**Supplementary Figure 1.**
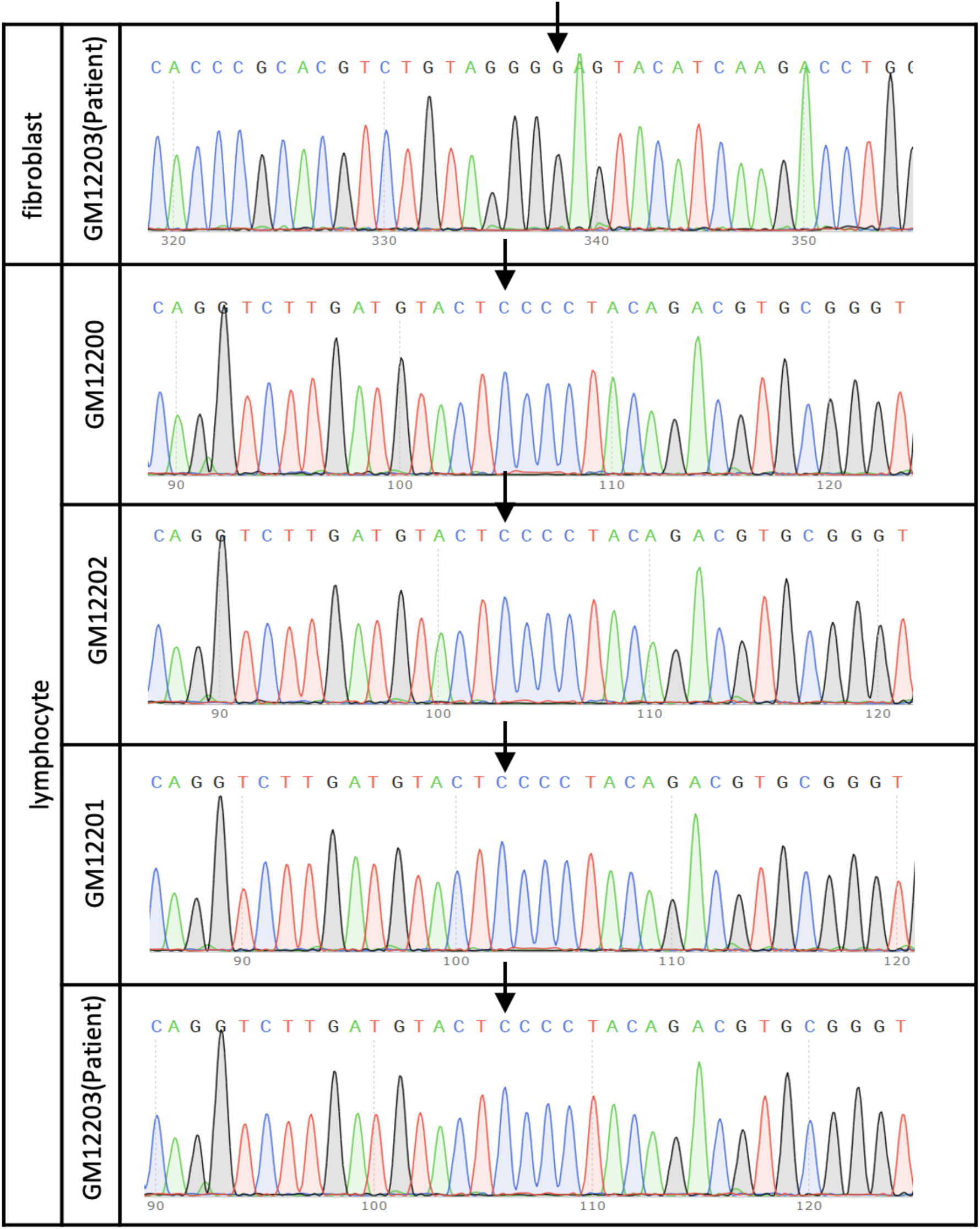
Sanger sequencing of Proteus syndrome patient fibroblasts, Proteus syndrome patient lymphocytes, and paired healthy lymphocytes. Arrows indicate the supposed location of the mutation.

**Supplementary Figure 2.**
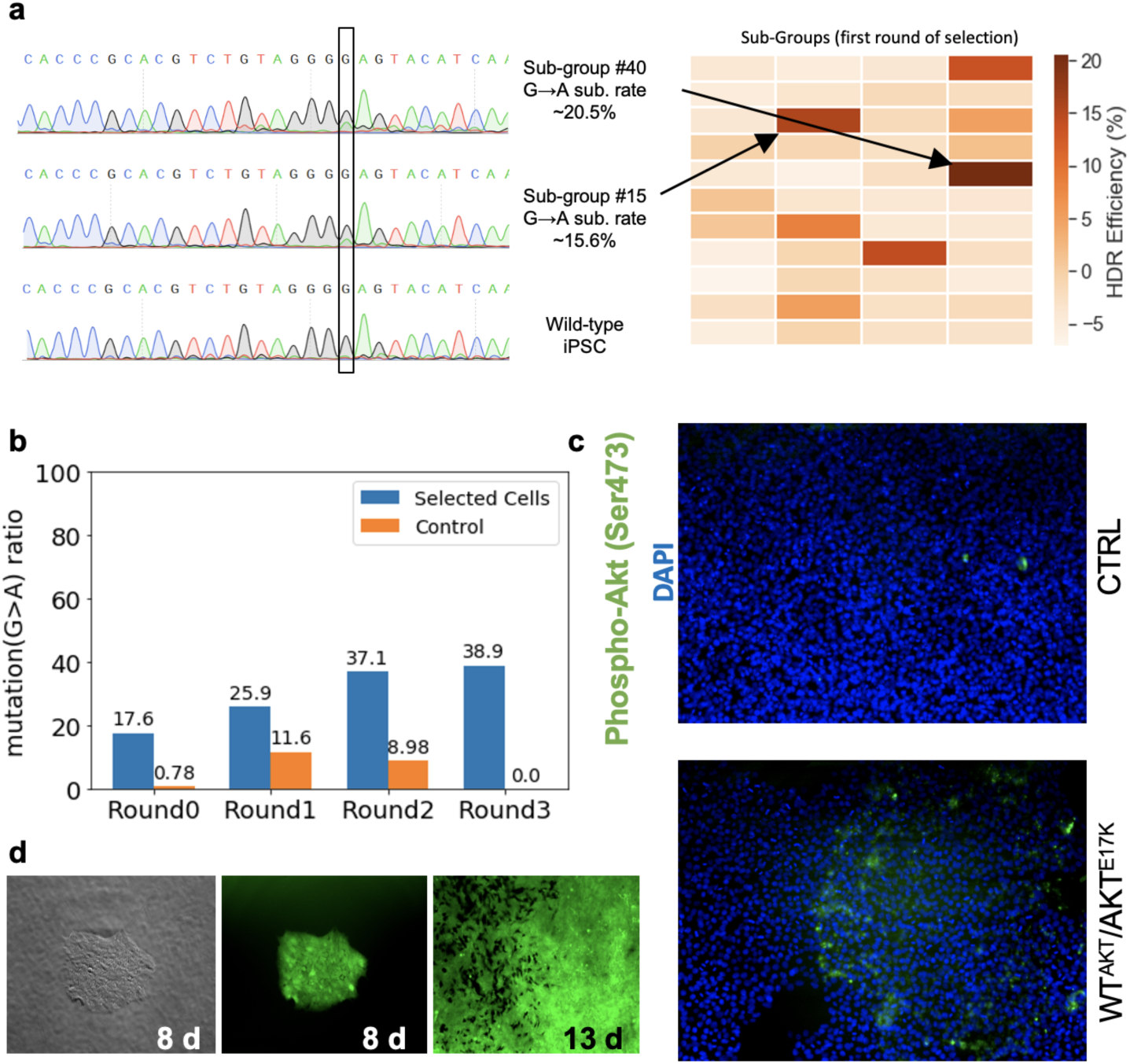
Analysis of CRISPR/Cas9 editing and lentiviral transduction. a) Sanger sequencing results for CRISPR/Cas9-edited iPSCs in 96-well plates. The cells were sorted into 96-well plates after editing, and each well included 10 cells to increase the chances of including viable cells for clone formation. The heatmap shows the HDR editing efficiency in 44 of 96 wells, while the cells in other wells did not survive. b) Mutation ratios (G→A) for different rounds of selection of CRISPR/edited cells. c) Immunofluorescence imaging of iPSCs stained with a phosphorylated AKT (Ser473) antibody and DAPI. d) Bright field and immunofluorescence imaging of lentivirus-transfected iPSCs.

**Supplementary Figure 3.**
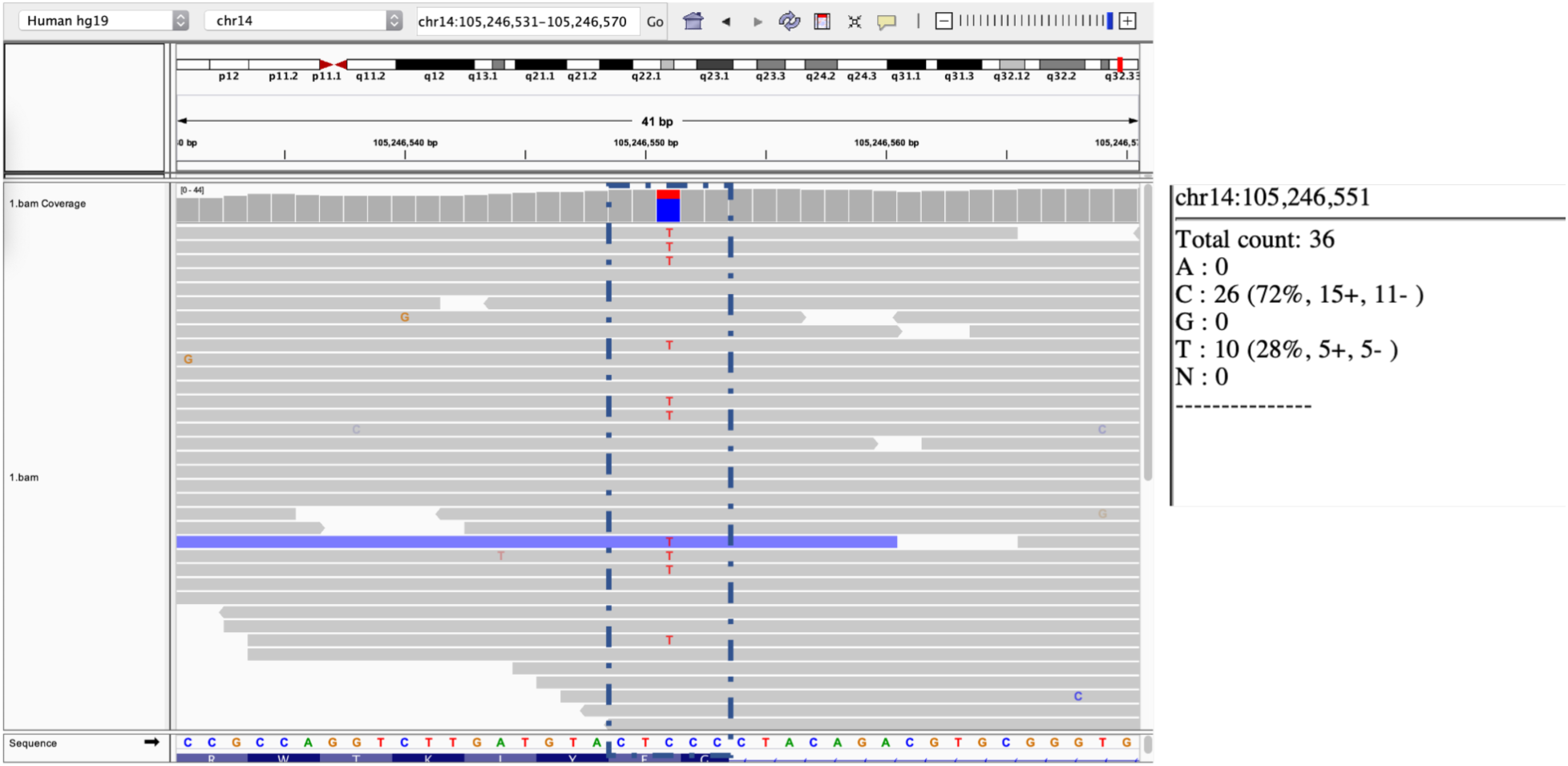
Whole-genome sequencing of purified CRISPR/Cas9-edited iPSCs.

**Supplementary Figure 4.**
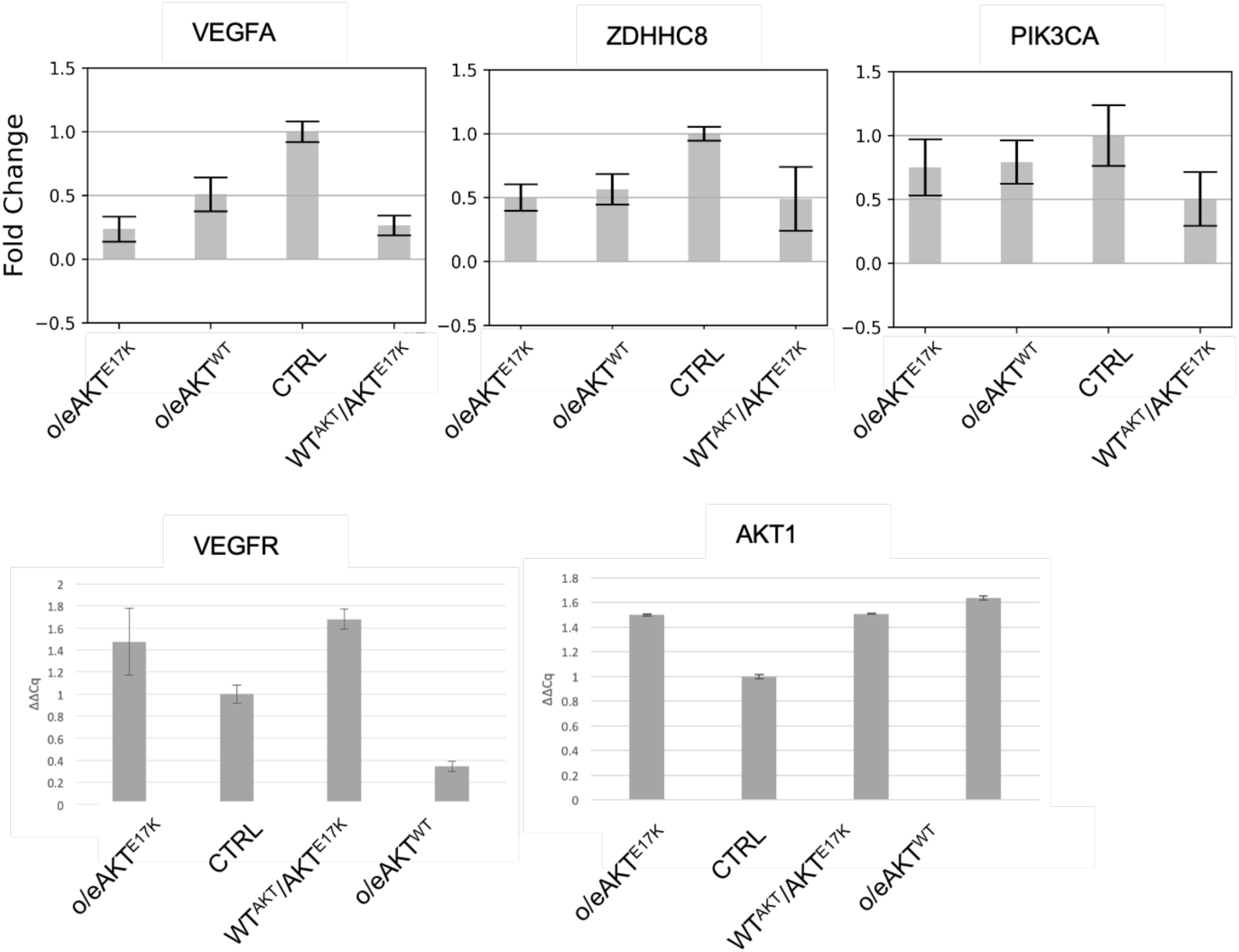
qPCR results for CRISPR/Cas9-edited or lentivirus-transfected iPSCs and control iPSCs.

**Supplementary Figure 5.**
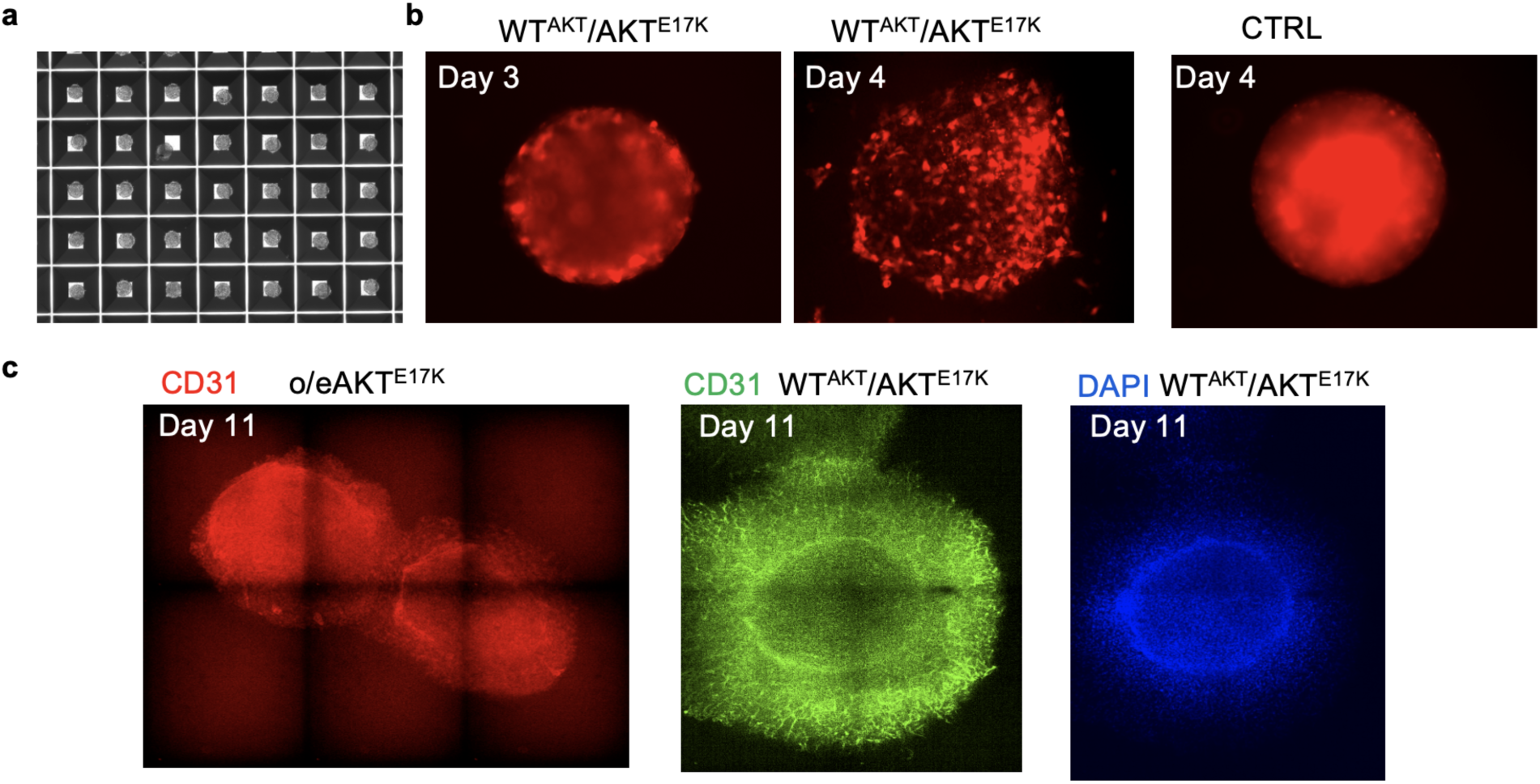
Bright-field imaging and confocal imaging of blood vessel organoids. a) Bright-field imaging of embryoid bodies on day 1 of organoid generation. b) Fluorescence imaging of RFP in WT^AKT^/AKT^E17K^ and CTRL organoids on day 3 and day 4. c) Confocal imaging of o/eAKT^E17K^ and WT^AKT^/AKT^E17K^ on day 11, after staining with a CD31 antibody and DAPI.

